# CD31^+^ T-cells express greater VEGF-A and CXCR4 levels than CD31^-^ counterparts with VEGF-A expression exacerbated with advancing age

**DOI:** 10.64898/2026.03.29.715132

**Authors:** Luke Stephen, Graham Wright, David J Muggeridge, Melanie Leggate, Vignesh Chandrakumar, Mark Ross

## Abstract

CD31^+^ T-cells reportedly possess angiogenic properties. These cells have recently been termed angiogenic T-cells (T_ANG_). Advancing age is associated with altered circulating T-cell phenotypes, including T_ANG_, and reduced angiogenesis. We examined various T_ANG_ subsets (CD3^+^, CD4^+^, CD8^+^), and their VEGF-A intracellular content in young (n=16, 18-30 years) and older (n=16, 50-65 years) male adults using flow cytometry. Cardiorespiratory fitness (*V̇*O_2_max) was quantified in all participants using a graded cycling ergometry test to volitional exhaustion. Resting blood samples were collected to measure circulating IL-6 and cytomegalovirus serostatus. CD31^+^ T-cells (T_ANG_) contained more VEGF-A than CD31^-^ T-cells (CD31^+^: 9374 ± 8587 AU vs CD31^-^: 8722 ± 8149 AU, *p* = 0.021) which was also exhibited in CD4^+^ and CD8^+^ subsets. Older adults possessed fewer CD4^+^ T_ANG_ cells as a proportion of total CD4^+^ T-cells than younger adults (young: 35 ± 11%; older: 24 ± 9%, *p* = 0.004), and CD3^+^ and CD4^+^ T_ANG_ subsets from older adults exhibited higher VEGF-A levels than younger adults (CD3^+^CD31^+^: young: 6081 ± 4001 AU; older: 13426 ± 10945 AU, *p* = 0.019; CD4^+^CD31^+^: young: 6373 ± 3972 AU; older: 15660 ± 12829 AU, *p* = 0.011). T_ANG_ cells were not associated with circulating IL-6, and T_ANG_ VEGF-A content was not associated with *V̇*O_2_max. Advancing age is associated with a pathological T_ANG_ phenotype, which may contribute to age-related inflammation and warrants further investigation as a potential therapeutic target.

## 1.0 Introduction

T-cells are a key part of the adaptive immune system, whose primary role is to combat infection (Sun *et al*., 2023). However, T-cells also demonstrate various non-immune functions such as tissue repair and maintenance (Arpaia *et al*., 2015). These adaptive immune cells are commonly found in secondary lymphoid organs as well as in the circulation. However, T-cells also reside in, and traffic to, various non-lymphoid organs such as the lungs, liver (Li *et al*., 2025), as well as skeletal muscle (Deyhle & Hyldahl, 2018), and therefore likely to contribute to local tissue maintenance.

A specific subset of T-cells which express the platelet endothelial cell adhesion molecule (PECAM; CD31) have been reported to possess angiogenic properties (Hur *et al*., 2007; Kushner *et al*., 2010a). These angiogenic T-cells (T_ANG_) have been shown to stimulate proliferation of endothelial progenitor cells (EPC) *in vitro* as well as promote perfusion recovery after hindlimb ischaemia in a mouse model (Hur *et al*., 2007) strongly suggesting that they act on endothelial cells to support vascular growth. These cells have strong migratory capacity, to both SDF-1 (commonly released from injured or ischaemic tissue) and VEGF-A (Kushner *et al*., 2010a). Together these observations suggest that these cells may play an important role in vascular tissue repair.

Ageing results in an altered immune system, characterised by inflammation, resulting in increased risk of autoimmune disorders and other chronic illnesses such as coronary artery disease and osteoporosis (Targonski *et al*., 2007). It is commonly acknowledged that primary indicators of an ageing immune system include a marked decline in T-cell proliferation (Han *et al*., 2023), and a shift from naïve to differentiated T-cell phenotype (Bruunsgaard *et al*., 1999; Soto-Heredero *et al*., 2023; Fraser & Owen, 2024). Together, these result in poor vaccine efficacy and adaptive immune response in older adults (Gustafson *et al*., 2020). There are several reports that T_ANG_ (defined as CD3^+^CD31^+^) cells are lower in older adults compared with younger counterparts (Ross *et al*., 2018a; Ross *et al*., 2018b). Moreover, these cells in older adults display a greater senescent profile (Ross *et al*., 2018a), reflective of a higher pro-inflammatory profile, which is characterised by an increased secretion of inflammatory cytokines IFN-γ, IL-6 and TNF-α (Zhang *et al*., 2021). In addition, the cell surface expression of a key chemokine receptor involved in cell migration and homing (CXCR4) is also reduced with advancing age (Ross *et al*., 2018b). These cells reportedly highly express the potent pro-angiogenic factor vascular endothelial growth factor-A (VEGF-A) (Hur *et al*., 2007), yet no study has investigated whether VEGF-A expression in T_ANG_ cells is affected by age, which may partly explain age-related impairment in angiogenic ability (Rivard *et al*., 1999).

Interestingly, a high cardiorespiratory fitness (CRF) has been shown to play a role in attenuating the negative effects of ageing on T-cells (Duggal *et al*., 2018), with individuals displaying higher CRF demonstrating a greater proportion of naïve T-cells and a decreased proportion of pro-inflammatory senescent T-cells (Spielmann *et al*., 2011). However, a key driver of immunological ageing and the associated immune phenotype shift is cytomegalovirus (CMV) serostatus. CMV infection is highly prevalent in older adults and is strongly associated with expansions of highly differentiated, senescent T-cells (Pawelec *et al*., 2009), as well as age-related immune dysfunction (Solana *et al*., 2012). Despite the established role of CMV in T-cell immunosenescence, no study to-date has investigated whether age, cardiorespiratory fitness, and/or CMV serostatus influences specific T_ANG_ subsets (CD4⁺ vs. CD8⁺), CXCR4⁺ cell numbers, or intracellular VEGF-A content. Therefore, this study investigated the effect of age, CRF, CMV serostatus and IL-6 concentrations on total CD3⁺CD31⁺ T-cells, CD4⁺ and CD8⁺ subsets, CXCR4 expression, and intracellular VEGF-A expression in humans.

## 2.0 Materials and Methods

### 2.1 Ethical Approval

The study was approved by Edinburgh Napier University’s Research Ethics and Governance Committee (reference number; 2842425).

### 2.2 Participants and Screening

Thirty-two healthy, non-obese (BMI<30kg·m^2^), non-smoking male participants aged 18-30 yrs (n=16) and 50-65 yrs (n=16) were recruited for this study. All participants gave written informed consent and completed health questionnaires to highlight any general health and cardiovascular concerns before data collection. Participants reported to the Edinburgh Napier University Human Performance Laboratory after an overnight fast, having not exercised for ≥24 h before the visit, having refrained from alcohol consumption the night before, and having not ingested caffeine the morning of the visit. Participants were measured for height and body mass (from this, body mass index (BMI) was calculated). Blood pressure (BP) was measured twice using an automated BP cuff (OMRON HEM-907, OMRON, Japan) after a 5-min supine rest. The average of the two blood pressure readings was then recorded.

### 2.3 Assessment of Cardiorespiratory Fitness

An incremental ramp exercise test to volitional exhaustion was performed to quantify maximum oxygen consumption (*V̇*O_2_max). The exercise test was performed on a cycle ergometer (Lode Corival, Lode B.V., Netherlands) with simultaneous breath- by-breath gas analysis (Cortex Metalyzer, Germany). Participants performed a warmup for 3-minutes, cycling at an initial power output of 100W for the younger participants and 50W for the older participants at 60–80 rpm. Following the warm-up, resistance was increase by 25W every minute until the participant could no longer maintain 60 rpm (Kim et al., 2016) or until volitional exhaustion. Heart rate was monitored throughout via heart telemetry (Polar H7 Heart Rate Monitors, Polar, Finland). *V̇*O_2_max was determined as the average *V̇*O_2_ of the last 30 seconds of the exercise test.

Participant characteristics are shown in **Table 1**.

**Table 1.** Participant Characteristics (n=32).

|  | 18-30 years (n=16) | 50-65 years (n=16) | *p* - value |
| --- | --- | --- | --- |
| Age (years) | 24 ± 2 | 57 ± 4 | <0.001* |
| Height (cm) | 183 ± 7 | 177 ± 7 | 0.010* |
| Body Mass (kg) | 83 ± 9 | 84 ± 12 | 0.668 |
| Body Mass Index (kg·m<sup>2</sup>) | 24 ± 2 | 26 ± 3 | 0.030* |
| Systolic Blood Pressure (mmHg) | 125 ± 8 | 131 ± 16 | 0.249 |
| Diastolic Blood Pressure (mmHg) | 81 ± 5 | 79 ± 8 | 0.500 |
| $\dot{V}O_{2\max}$ (ml·kg·min<sup>-1</sup>) | 39 ± 7 | 29 ± 10 | 0.006* |

Values shown are mean ± SD. \**p* < 0.05 between age groups.

### 2.4 Blood Collection

Venepuncture was performed with the participants in a seated, upright position after a 5-minute rest period, prior to the assessment of *V̇*O_2_max. A 21-guage needle and collection kit (BD Biosciences, USA) were used for collecting the blood samples. Blood samples were collected into 6 x 10ml EDTA coated tubes and 1 x 6ml serum separating tube using the BD Vacutainer Safety-Lok system (BD Biosciences, USA). Blood samples were used for T-cell analysis (see Peripheral Blood Mononuclear Cell Separation and Flow Cytometric Quantification of CD31^+^ T-Cells), serum analysis for cytomegalovirus (CMV) and plasma analysis for IL-6 concentrations (see Plasma and Serum Analysis for IL-6 and CMV Serostatus)

### 2.5 Peripheral Blood Mononuclear Cell Separation and Flow Cytometric Quantification of CD31^+^ T-Cells

Peripheral blood mononuclear cells (PBMC) were isolated with a density gradient centrifugation using Lymphoprep^TM^ (Axis-Shield, USA) as performed previously (Ross *et al*., 2018b).

For flow cytometric enumeration of CD31^+^ T-cells, firstly, 1.0 x 10^6^ mononuclear cells were incubated with monoclonal antibodies targeted to cell surface markers (2μL of anti-CD3-FITC, anti-CD4- BV510, anti-CD8-PE-Cy5, anti-CD31-BV421 and anti-CXCR4-PE-Cy7 - all Biolegend, USA) for 30 minutes at 4°C in the dark. The cells were then permeabilised using a permeabilization buffer (BD Cytoperm, BD Biosciences, USA) and incubated for 20 minutes at 4°C in the dark. The cells were then washed using a wash buffer (BD Perm/Wash, BD Biosciences, USA) followed by incubation with 2μL anti-VEGF-A (Abcam, UK) for 30 minutes at 4°C in the dark. Immediately before flow cytometric enumeration, 500μL PBS-BSA was added. T_ANG_ cells (defined as: CD3^+^CD31^+^, CD3^+^CD4^+^CD31^+^, CD3^+^CD8^+^CD31^+^) CXCR4 cell surface expression and intracellular VEGF-A content were quantified on a 3-laser flow cytometer (BD FACSCelesta, BD Biosciences). A minimum of 20,000 lymphocyte events were collected per sample. Isotypes for both CXCR4-PE-Cy7 (Biolegend, USA) and VEGF-A-PE (Abcam, UK) were used in matched concentrations as controls to distinguish between positive and negative events, and fully stained non-permeabilised isotype controls were also used. Compensation was applied to correct for fluorescent spillover between detectors and channels using appropriate compensation beads (BD™ CompBeads, BD Biosciences, USA) and single stained controls.

After data acquisition, data were analysed using Flow Logic (FlowLogic Version 8.7, FlowLogic, Australia). The percentage of lymphocyte subsets of interest were analysed (CD3^+^CD31^+^ cells, with co-expression of CXCR4, as well as CD4^+^ and CD8^+^ subsets) and circulating number were calculated by multiplying the percentage count as quantified by semi-automated haematology analyser (XS-1000i, Sysmex, Japan). All haematology analyser data were measured in duplicate and averaged. Expression of VEGF-A within these subsets was expressed as median fluorescent intensity (MFI).

Flow cytometry gating strategy is shown in **Figure 1**.

**Figure 1.**
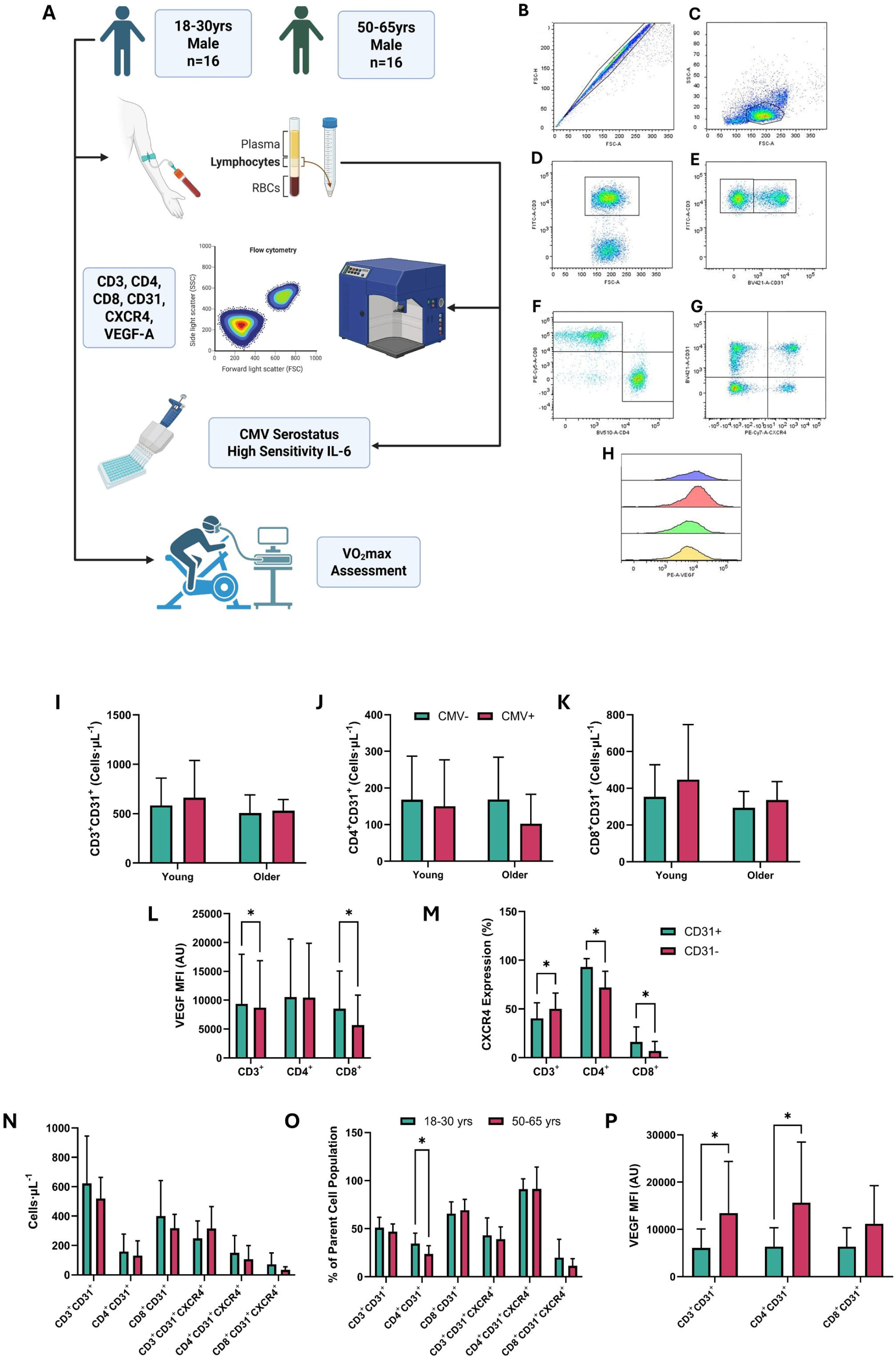
Study Design, Gating Strategies for the Flow Cytometric Quantification of T_ANG_ Cells and Differences in T_ANG_ Cells and VEGF and CXCR4 Expression by Phenotype and Age. *A: Young (18-30 years) and older (50-65 years) adults provided blood samples for quantification of T-cells, IL-6 and CMV serostatus, and underwent a graded exercise test to exhaustion for quantifying V̇O_2_max. B-G: Gating strategy for T_ANG_ cell quantification: first, identification of singlet cells (B), subsequent detection of the lymphocyte population, forward scatter vs. side scatter (FSC vs. SSC; C), and identification of the CD3^+^ T-cell population (D) and the CD31^+^ and CD31^-^ subsets (E), and further gating of CD4^+^ population and CD8^+^ populations (F). Finally, these expression of CXCR4 (G) and VEGF (H) were quantified in these cells. I-K: No effect of CMV serostatus on CD3^+^CD31^+^ (I), CD4^+^CD31^+^ (J) or CD8^+^CD31^+^ (K) T-cells. L-N: Analysis for differences in VEGF (L) and CXCR4 (M) between CD31^+^ and CD31^-^ T-cells. N-P: Differences in T_ANG_ subsets (N + O) and T_ANG_ VEGF expression (P) between age groups. Values shown are mean ± SD, *p < 0.05*

### 2.6 Plasma and Serum Analysis for IL-6 and CMV Serostatus

After phlebotomy, serum vacutainers were left at room temperature for a minimum of 30 minutes prior to centrifugation. The vacutainers were subsequently centrifuged at 900 x *g* for 15 min at 4 °C, with serum aliquoted and stored at -80°C until analysis. Serum and plasma samples were thawed and used to quantify CMV serostatus (positive, negative) and IL-6 concentrations (pg·mL^-1^), respectively, using commercially available enzyme-linked immunosorbent assays (ELISA; Human Anti-CMV IgG ELISA Kit, Abcam, Cambridge, UK; catalogue number ab108724; Human IL-6 Quantikine High Sensitivity ELISA Kit, R&D Systems, Minneapolis, MN, USA; catalogue number HS600C). Analysis was performed for both assays using a plate reader (Multiskan FC, Thermofisher, USA), with fluorescence assessed using a 450nm filter. Samples were analysed in duplicates, with intra-assay coefficient of variation of 5.5% and 6.6% for the IL-6 and CMV assay, respectively. The lower limit of detection for the IL-6 assay was reported as 0.03 pg·mL^-1^.

### 2.7 Statistical Analysis

A Shapiro-Wilk test was used to confirm that the data were normally distributed. Paired T-tests were performed to assess VEGF-A MFI and CXCR4 expression differences between CD31^+^ and CD31^-^ T-cells. Subsequently, independent T-tests were performed to assess whether there were any significant differences in circulating T_ANG_ number (CD3^+^CD31^+^, CD4^+^CD31^+^, CD8^+^CD31^+^, and CXCR4^+^ subsets) and T_ANG_ intracellular VEGF-A expression between young and older adults. Multiple linear regressions were performed to determine the relationship between age and cardiorespiratory fitness and various T_ANG_ subsets, expression of cell surface CXCR4 and intracellular VEGF-A. Data were analysed using GraphPad Prism (version 9, Dotmatics, USA) and Jamovi (version 2.7.17; https://www.jamovi.org). Significance was set at *p* < 0.05. Data are presented as means ± SD unless stated otherwise.

## 3.0 Results

### 3.1 CD31^+^ T-cells are not influenced by CMV serostatus

Comparisons between CMV seropositive and seronegative individuals demonstrate that there were no statistically significant differences between any CD3^+^CD31^+^ T-cell subset (proportion or absolute counts), T_ANG_ CXCR4 or VEGF-A expression (all *p* > 0.05; absolute T_ANG_ data shown in **Figure 1**).

### 3.2 CD31^+^ T-cells express greater intracellular VEGF content than CD31^-^ T-cells

Our analysis demonstrates that CD31^+^ T-cells contain greater intracellular VEGF-A content (defined as VEGF MFI) than CD31^-^ counterparts. This finding persists for total CD31^+^ T-cells (CD31^+^: 9374 ± 8587 AU vs CD31^-^: 8722 ± 8149 AU, *p* = 0.021) and CD8^+^CD31^+^ T-cells (CD31^+^: 8523 ± 6528 AU vs CD31^-^: 5667 ± 5206 AU, *p* < 0.001). However, no statistically significant difference was found for CD4^+^CD31^+^ T-cells (*p* = 0.791) CXCR4 expression was greater in CD31^-^ vs CD31^+^ T-cells (CD31^+^: 41 ± 16%; CD31^-^: 49 ± 16%, *p* = 0.011). However, upon further analysis of the CD4^+^ and CD8^+^ subsets, CXCR4 expression was greater in CD4^+^CD31^+^ vs CD4^+^CD31^-^ cells (93 ± 8%; 72 ± 17%, *p* < 0.001) and similar was observed in the CD8^+^ subset (16 ± 16%; 7 ± 10%, *p* < 0.001).

CXCR4 expression was significantly higher in CD4^+^CD31^+^ T-cells than CD8^+^CD31^+^ T cells (CD4^+^CD31^+^: 88 ± 20%; CD8^+^CD31^+^: 16 ± 16%, *p* < 0.001). VEGF-A expression was not different between CD4^+^CD31^+^ and CD8^+^CD31^+^ T-cells (*p* = 0.183).

Data is shown in **Figure 1**.

### 3.3 Older adults display reduced proportion of CD4^+^ T-Cells expressing CD31 than younger adults

There were no significant differences in CD3^+^CD31^+^ or CD8^+^CD31^+^ T-cells between young and older adults, either as absolute counts, or proportion data (all *p* > 0.05). Older adults did display fewer CD4^+^CD31^+^ cell profile than younger adults, represented as a proportion of all T-cells (young: 35 ± 11%; older: 24 ± 9%, *p* = 0.004) but not as absolute counts (young: 159 ± 119 cells·μL^-1^; older: 131 ± 100 cells·μL^-1^, *p* = 0.482).

There were no differences in CXCR4^+^ T_ANG_ counts between the age groups (all *p* <0.05), however T_ANG_ VEGF-A MFI was significantly higher in older adults compared to younger adults in CD3^+^CD31^+^ and CD3^+^CD4^+^CD31^+^ subsets (CD3^+^CD31^+^: young: 6081 ± 4001 AU; older: 13426 ± 10945 AU, *p* = 0.019; CD4^+^CD31^+^: young: 6373 ± 3972 AU; older: 15660 ± 12829 AU, *p* = 0.011).

Data for simple age comparisons is shown in **Table 2**.

**Table 2.** Angiogenic T-Cell Number and Expression of CXCR4 and VEGF in Young (18-30 years) and Older (50-65 years) Adults (n=32).

|  | 18-30 years<br>(n=16) | 50-65 years<br>(n=16) | *p* - value | *t*-statistic |
| --- | --- | --- | --- | --- |
| **CD3<sup>+</sup>CD31<sup>+</sup>** |  |  |  |  |
| **Cells·μL<sup>-1</sup>** | 623 ± 322 | 519 ± 144 | 0.250 | 1.174 |
| **Proportion (%)** | 51 ± 11 | 47 ± 8 | 0.230 | 1.225 |
| **CXCR4<sup>+</sup> Cells·μL<sup>-1</sup>** | 248 ± 119 | 315 ± 149 | 0.167 | 1.425 |
| **Proportion (%)** | 43 ± 18 | 39 ± 13 | 0.480 | 0.715 |
| **VEGF MFI** | **6081 ± 4001** | **13426 ± 10945** | **0.019\*** | **2.496** |
| **CD4<sup>+</sup>CD31<sup>+</sup>** |  |  |  |  |
| **Cells·μL<sup>-1</sup>** | 159 ± 119 | 131 ± 100 | 0.482 | 0.712 |
| **Proportion (%)** | **35 ± 11** | **24 ± 9** | **0.004\*** | **3.135** |
| **CXCR4<sup>+</sup> Cells·μL<sup>-1</sup>** | 149 ± 118 | 120 ± 97 | 0.481 | 0.715 |
| **Proportion (%)** | 91 ± 11 | 96 ± 3 | 0.135 | 1.539 |
| **VEGF MFI** | **6373 ± 3972** | **15660 ± 12829** | **0.011\*** | **2.748** |
| **CD8<sup>+</sup>CD31<sup>+</sup>** |  |  |  |  |
| **Cells·μL<sup>-1</sup>** | 400 ± 242 | 318 ± 95 | 0.215 | 1.266 |
| **Proportion (%)** | 62 ± 13 | 68 ± 11 | 0.386 | 0.880 |
| **CXCR4<sup>+</sup> Cells·μL<sup>-1</sup>** | 37 ± 38 | 28 ± 19 | 0.061 | 1.950 |
| **Proportion (%)** | 20 ± 19 | 11 ± 7 | 0.109 | 1.653 |
| **VEGF MFI** | 6660 ± 3946 | 11216 ± 8056 | 0.063 | 1.942 |

*Values shown are mean ± SD, MFI- Mean Fluorescence Intensity. \* *p* <0.05 between age groups.*

### 3.4 CD8+CD31^+^ T-cells influenced by cardiorespiratory fitness, independent of age

The multiple linear regression analyses demonstrated that CD4^+^CD31^+^ T_ANG_ (% of CD4^+^were inversely associated with age (*r*^2^ = 0.148, *F* = 5.060, *p* = 0.032), with no other subset demonstrating a statistically significant relationship. After controlling for age, CRF was inversely associated with CD8^+^CD31^+^ T_ANG_ (% of CD8^+^) (*r*^2^ = 0.201, *F* = 0.750, *p* = 0.011). No other relationships between cardiorespiratory fitness and T_ANG_ subset were observed (all *p* > 0.05).

For other subsets, for both simple, and multiple regression (control for age) revealed no association between circulating T_ANG_ CXCR4 or VEGF-A expression and cardiorespiratory fitness levels (*V̇*O_2_max). Summary statistics are displayed in **Table 3**.

**Table 3.** Linear Regression between CD31^+^ T-Cell Number and Cardiorespiratory Fitness (n=32).

| T<sub>ANG</sub> Subset | Model | R<sup>2</sup> | F - Value | p - Value | R<sup>2</sup> Change | F-Change | Change p - Value |
| --- | --- | --- | --- | --- | --- | --- | --- |
| CD3<sup>+</sup>CD31<sup>+</sup> | 1 | 0.030 | 0.881 | 0.356 | - | - | - |
|  | 2 | 0.075 | 1.132 | 0.337 | 0.045 | 1.370 | 0.251 |
| CD3<sup>+</sup>CD31<sup>+</sup> (% of CD3<sup>+</sup>) | 1 | 0.022 | 0.664 | 0.664 | - | - | - |
|  | 2 | 0.120 | 1.905 | 0.168 | 0.097 | 3.100 | 0.089 |
| CD4<sup>+</sup>CD31<sup>+</sup> | 1 | 0.001 | 0.020 | 0.888 | - | - | - |
|  | 2 | 0.004 | 0.052 | 0.949 | 0.003 | 0.085 | 0.773 |
| CD4<sup>+</sup>CD31<sup>+</sup> (% of CD4<sup>+</sup>) | 1 | **0.148** | **5.060** | **0.032\*** | - | - | - |
|  | 2 | 0.153 | 2.53 | 0.097 | 0.005 | 0.158 | 0.694 |
| CD8<sup>+</sup>CD31<sup>+</sup> | 1 | 0.055 | 1.690 | 0.204 | - | - | - |
|  | 2 | 0.107 | 1.670 | 0.207 | 0.051 | 1.610 | 0.215 |
| CD8<sup>+</sup>CD31<sup>+</sup> (% of CD8<sup>+</sup>) | 1 | 0.047 | 1.440 | 0.239 | - | - | - |
|  | 2 | **0.249** | **4.630** | **0.018\*** | **0.201** | **7.500** | **0.011\*** |

*Model 1- Age* *Model 2- Age and $\dot{V}O_{2max}$* *\*p < 0.05.*

**Table 4.** Linear Regression between CD31^+^ T-Cell CXCR4, VEGF-A Expression and Cardiorespiratory Fitness (n=32).

| T<sub>ANG</sub> Subset | Model | R<sup>2</sup> | F - Value | p - Value | R<sup>2</sup> Change | F - Change | Change p - Value |
| --- | --- | --- | --- | --- | --- | --- | --- |
| CD3<sup>+</sup>CD31<sup>+</sup>CXCR4<sup>+</sup> (%) | 1 | 0.005 | 0.133 | 0.718 | - | - | - |
|  | 2 | 0.136 | 2.209 | 0.129 | **0.132** | **4.270** | **0.048\*** |
| CD3<sup>+</sup>CD31<sup>+</sup> VEGF MFI | 1 | **0.173** | **5.640** | **0.025\*** | - | - | - |
|  | 2 | 0.186 | 2.960 | 0.069 | 0.013 | 0.411 | 0.527 |
| CD4<sup>+</sup>CD31<sup>+</sup>CXCR4<sup>+</sup> (%) | 1 | 0.110 | 3.320 | 0.080 | - | - | - |
|  | 2 | 0.156 | 2.410 | 0.110 | 0.047 | 1.45 | 0.240 |
| CD4<sup>+</sup>CD31<sup>+</sup> VEGF MFI | 1 | **0.204** | **6.900** | **0.014\*** | - | - | - |
|  | 2 | **0.212** | **3.500** | **0.045\*** | 0.008 | 0.273 | 0.606 |
| CD8<sup>+</sup>CD31<sup>+</sup>CXCR4<sup>+</sup> (%) | 1 | 0.128 | 3.950 | 0.057 | - | - | - |
|  | 2 | 0.145 | 2.200 | 0.131 | 0.017 | 0.523 | 0.476 |
| CD8<sup>+</sup>CD31<sup>+</sup> VEGF MFI | 1 | 0.159 | 3.590 | 0.073* | - | - | - |
|  | 2 | 0.168 | 1.810 | 0.192 | 0.009 | 0.184 | 0.673 |

*Model 1- Age* *Model 2- Age and $\dot{V}O_{2max}$* *\*p < 0.05. T<sub>ANG</sub> CXCR4 data presented as proportion data only. MFI- Median Fluorescence Intensity.*

### 3.5 Circulating CD31^+^ T cells are not associated with serum IL-6 concentrations in young and older adults

Older adults demonstrated a higher circulating level of IL-6 than younger individuals (young: 0.322 ± 0.142 pg·mL^-1^; older: 0.643 ± 0.421 pg·mL^-1^, *p* = 0.025). Linear regression analysis revealed for both simple and multiple regressions (control for age) revealed no association between circulating IL-6 levels and other T_ANG_ subsets, T_ANG_ CXCR4 or VEGF-A expression (all *p* > 0.05).

## 4.0 Discussion

The study’s main findings are that CD31^+^ T-cells express greater VEGF-A, a key angiogenic growth factor, than CD31^-^ T-cells. Paradoxically, whilst CD31^+^ T-cell numbers did not differ between age groups, VEGF-A expression was elevated in in CD31^+^ T-cells from older adults, suggesting a higher basal inflammatory profile. In addition, we demonstrated that these angiogenic T-cells also have greater CXCR4 expression than CD31^-^ T-cells, suggestive of a greater migratory function to SDF-1, which is elevated in tissue injury (Ratajczak *et al*., 2004).

VEGF-A is a highly pro-angiogenic growth factor (Wagner, 2011), linked to both hypoxia-and exercise-induced angiogenesis (Wagner, 2011; Thom *et al*., 2014; Ross *et al*., 2023). Various cell types produce and secrete VEGF (Stepanova *et al*., 2007) including neutrophils (Gong & Koh, 2010), monocytes/macrophages (Stepanova *et al*., 2007), endothelial cells (Guangqi *et al*., 2012), and skeletal muscle cells (Ryan *et al*., 2006; Delavar *et al*., 2014). In the 1980s, several studies demonstrated that T-cells produce VEGF, known then as vascular permeability factor (Tomizawa *et al*., 1985; Heslan *et al*., 1986). In 2007, a study by Hur *et al*. (2007) demonstrated that a specific subset of T-cells that express CD31, were required for optimal *in vitro* growth of endothelial progenitor cells, with subsequent studies demonstrating a pro-angiogenic phenotype of these CD31^+^ T-cells, with work by Kushner *et al*. (2010a) observing greater migration to VEGF and SDF-1 by CD31^+^ vs CD31^-^ T-cells. Our study further emphasises that these CD31^+^ T-cells likely possess greater angiogenic function than CD31^-^ T-cells through enhanced VEGF-A production, and greater CXCR4 expression albeit in CD4^+^ and CD8^+^ subsets only. Interestingly, we observed lower CXCR4 expression in CD3^+^CD31^+^ T-cells as the parent population, likely driven by non-CD4^+^ or CD8^+^ cells such as γδ T-cells (Wang *et al*., 2025), however, these were not quantified in this study, and future work should include analysis of CD31^+^ expression and pro-angiogenic activity of such cells, as these cells are good candidates for cellular therapies, due to the lower risk of graft versus host disease (Revesz *et al*., 2024).

Older adults display a lower number of circulating CD31^+^ T-cells compared to younger adults (Hur *et al*., 2007; Kushner *et al*., 2010b; Ross *et al*., 2018a; Ross *et al*., 2018b), with these cells also demonstrating impaired migration, increased apoptosis susceptibility, reduced telomerase activity (Kushner *et al*., 2010b), and greater senescence (Ross *et al*., 2018a) compared to younger adults. Together, these findings suggest impaired angiogenic activity of these putative angiogenic T-cells (T_ANG_), however, our data in this study show that T_ANG_ cells from older adults possess greater VEGF-A content than younger adults. Rather than this indicating a greater angiogenic capability in these T-cells, we suggest this excess unstimulated VEGF-A production by these cells in older adults is indicative of a chronic inflammatory state. Indeed it is known that excessive VEGF-A signalling is associated with several inflammatory states, including diabetic retinopathy (Gu *et al*., 2020; Ahmad & Nawaz, 2022), cancer (Freeman *et al*., 1995; Ahmad & Nawaz, 2022) and rheumatoid arthritis (Foster *et al*., 2009). There is evidence of excess VEGF-A production by T-cells (CD4^+^) in type 1 diabetics (Marek *et al*., 2010), as well as in patients with chronic obstructive pulmonary disease (Mikko *et al*., 2009), which may be a key contributor to tissue inflammation in these states. The data in our study suggests that whilst CD31^+^ T-cells contain more VEGF-A than CD31^-^ T-cells, which likely contributes to the vascular phenotype of these T-cells, may also, with advancing age, contribute to T-cell mediated inflammation in age-related disease (Macaulay *et al*., 2013; Yu *et al*., 2016; Bektas *et al*., 2017). We did not co-stain for CD28 to perform sub-analysis on VEGF-A production in CD28^null^, senescent-associated, T_ANG_, with these being elevated in older adults (Ross *et al*., 2018a) and themselves containing more intracellular pro-inflammatory cytokines (IL-6, TNF- and IFN-γ) compared to CD28^+^ T_ANG_ cells (Zhang *et al*., 2021), may be the source of the excess VEGF-A, rather than all T_ANG_ cells per se. Therefore, as a result of our data and those presented in this discussion, the higher VEGF-A content of T_ANG_ cells from older adults presents a propensity for pathological angiogenesis and chronic inflammation, and future studies are needed to identify source of the elevated VEGF-A, and whether this is associated or contributes to inflammatory conditions.

Our subsequent analyses investigated whether cardiorespiratory fitness was associated with our T_ANG_ measures. Cardiorespiratory fitness, which is associated with altered T-cell phenotypes in several reports (Spielmann *et al*., 2011) had limited influence on these T_ANG_ cells, with only CD8^+^ T_ANG_ cells demonstrating a significant relationship with cardiorespiratory fitness. Initially there was a negative relationship between CD8^+^ T_ANG_ cells and *V̇*O_2_max, which is surprising, given CD31 is a marker of recent thymic emigrants, and previous work has demonstrated that *V̇*O_2_max is positively associated with naïve CD8^+^ T-cells (Spielmann *et al*., 2011), which are expected to be more recent emigrants from thymic development than differentiated, effector T-cells. However, after correcting for age, as our older adults possessed significantly lower levels than their younger counterparts, and previous evidence that ageing is associated with a decline in *V̇*O_2_max (Rogers *et al*., 1990), there was no significant relationship, indicating that fitness per se, did not influence this subset. Interestingly, cardiorespiratory fitness was positively associated with CD8^+^ T_ANG_ CXCR4 expression which remained significant after controlling for age, suggesting that regular exercise, or physical activity, may influence T_ANG_ migratory potential. We did not control for CMV serostatus, as our analysis demonstrated that cell number and various measures of T_ANG_ subsets were not influenced by CMV serostatus. Our data also shows that older adults displayed a higher BMI than younger adults, and so may be a confounding variable in our analysis. However, after controlling for age, BMI was not associated with any T_ANG_ subset or measure. Further studies are required to elucidate the direct effects of exercise and physical activity interventions on these cells, using *in vitro* and *in vivo* studies to determine the potential of these cells to migrate to tissue injury.

Interestingly, our T_ANG_ cellular data was not associated with plasma levels of the pro-inflammatory cytokine IL-6, which is in contrast to previous published works (Zhang *et al*., 2021). The primary reason for this discrepancy between these studies is the lack of CD28 phenotyping of our T_ANG_ cells, as Zhang *et al*. (2021) observed a positive relationship between circulating IL-6 and CD28^null^CD4^+^ T_ANG_ cells, but did not present data on the parent population (e.g. CD4^+^ T_ANG_). The senescent subset of T_ANG_ cells produce more proinflammatory cytokines (IL-6, TNF- IFN-γ) (Zhang *et al*., 2021) which may contribute to elevated circulating inflammatory milieu with advancing age, however, we cannot determine this from our data, which is a limitation of our study. Also, IL-6 was the only pro-inflammatory cytokine measured in this study, and so other age-related inflammatory factors, such as C-reactive protein (CRP) and TNF-α, may, in future, provide more clarity on relationship between immune cell subsets, age and inflammation. It is clear that a more in-depth phenotyping strategy is required to fully elucidate the impact of inflammation, age, exercise, cardiorespiratory fitness on these T-cells.

### 4.1 Limitations

As noted above, this study is not limitations. Firstly, the flow cytometric phenotypic strategy for this study omitted CD28, thus preventing our ability to determine VEGF-A content between senescent and non-senescent T_ANG_ cells to better understand the relationship between these cells and advancing age. Secondly, the participant pool was wholly male, and therefore the findings cannot be generalisable to female populations. Future studies are planned to examine how ageing and menopause influence T_ANG_ populations, which may yield deeper mechanistic insights into the relationship between menopausal transition and vascular inflammation.

### 4.2 Conclusion

This study is the first to demonstrate that CD31^+^ putative angiogenic T-cells (T_ANG_) possess greater VEGF-A content and CXCR4 expression than CD31^-^ T-cells, suggestive of a greater angiogenic capacity. However, older adults display an excessive T_ANG_ VEGF-A profile, which is likely to reflect a shift from physiological to pathological angiogenesis and inflammation with advancing age. Cardiorespiratory fitness was not associated with T_ANG_ VEGF-A content but was positively associated with CD8^+^ T_ANG_ CXCR4 expression, indicating that T_ANG_ migratory capacity (a key function of such cells in angiogenesis) is higher in those with a higher *V̇*O_2_max. Overall, these findings show that ageing drives a distinct shift toward a more inflammatory and dysregulated T_ANG_ phenotype, marking these cells as sensitive indicators of immune–vascular ageing

### 4.3 Author Statement

Luke Stephen, Mark Ross and Graham Wright designed the study and conceived the research question. Luke Stephen, Mark Ross, Vignesh Chandrakumar and Graham Wright collected data. Luke Stephen and Mark Ross analysed the data. All authors contributed to the interpretation of the data for the work. Luke Stephen and Mark Ross drafted the manuscript. All authors revised the manuscript, approved the final version, and agree to be accountable for all aspects of the work. All authors had full access to and accept responsibility for the data presented in this report

